# Spindle F-actin coordinates the first metaphase-anaphase transition in *Drosophila* meiosis

**DOI:** 10.1101/2022.08.09.503402

**Authors:** Benjamin W. Wood, Timothy T. Weil

## Abstract

Meiosis is a highly conserved feature of sexual reproduction that ensures germ cells have the correct number of chromosomes prior to fertilization. A subset of microtubules, known as the spindle, are essential for accurate chromosome segregation during meiosis. Building evidence in mammalian systems has recently highlighted the unexpected requirement of the actin cytoskeleton in chromosome segregation; a network of spindle actin filaments appear to regulate many aspects of this process. Here we show that *Drosophila* oocytes also have a spindle population of actin that regulates the formation of the microtubule spindle and chromosomal movements throughout meiosis. We demonstrate that genetic and pharmacological disruption of the actin cytoskeleton has a significant impact on spindle morphology, dynamics, and chromosome alignment and segregation during the metaphase-anaphase transition. We further reveal the requirement of calcium in maintaining the microtubule spindle and spindle actin. Together, our data highlights the significant conservation of morphology and mechanism of the spindle actin during meiosis.

## INTRODUCTION

Meiosis is a well studied and documented process that is essential for the production of haploid gametes during sexual reproduction. In mammals, fetal oogonia initiate meiosis synchronously, arresting at prophase I and pausing in this state until sexual maturity, whereupon an oocyte or small subsets of oocytes are released periodically and meiosis is resumed^1^. Nuclear envelope breakdown initiates this resumption, resulting in the formation of a bipolar spindle network as microtubules polymerize, capture chromosomes and then align them on the metaphase plate^2^. The spindle interacts closely with the cytoplasmic actin, which aids in the asymmetric positioning of the spindle adjacent to the cortex, enabling asymmetric cell division to leave a single oocyte containing the necessary maternal components^2-7^. Oocytes are arrested in metaphase II until fertilization and subsequent egg activation. An intracellular calcium rise then triggers the completion of meiosis, resulting in the formation of the maternal pronucleus which undergoes fusion with the paternal pro-nucleus to form a diploid zygote^8,9^.

In *Drosophila melanogaster*, the process is similar but with a few key differences. Oocytes are generated continuously when environmental conditions are favorable. Connected to supporting cells via cytoplasmic bridges and surrounded by a mono-layer of epithelial cells, the oocyte passes through morphologically distinct stages during oogenesis. Meiosis is held in late prophase I for the majority of oogenesis, until the prophase-metaphase transition^10,11^. Key features of this step can be observed within the *Drosophila* oocyte, such as nuclear envelope breakdown and formation of a bipolar spindle as microtubule spindles capture the meiosis I chromosomes^12,13^.

Unlike mammals and most vertebrates, the final arrest in *Drosophila* oocytes is at metaphase I, and the bipolar spindle structure that forms is more defined and has more focused spindle poles when compared to the “barrel” shaped spindles of mammals. Activation in *Drosophila* occurs prior to fertilization as the oocyte passes into the oviduct, resulting in calcium influx through transient receptor potential melastatin (TrpM) ion channels in the plasma membrane of the oocyte^14,15^. This calcium event enables the resumption of meiosis from its arrested state, and can be observed using a variety of microtubule labelling tools^16^. Detailed cytological studies of *Drosophila* oocytes revealed that at the metaphase I arrest, the chromosomes exists as a central mass, including the non-exchange chromosomes, which can be visible as a separate entity during pro-metaphase or anaphase^16,17^. The metaphase to anaphase transition is described as a stereotypical series of events; 1) the meiotic spindle elongates; 2) then contracts, decreasing in length but increasing in width; 3) the meiotic spindle then rotates in relation to the cortex resulting in a perpendicular orientation to the cortex^16^. Identification of *Drosophila* oocytes at metaphase I arrest is easily observable through the developmental stage of the dorsal appendages, which at a length of greater than 250 μm indicate the mature oocyte^17^.

The mature *Drosophila* oocyte itself is approximately 500 μm in length, compared to the spindle which is approximately 10 μm (∼50:1 ratio of oocyte to spindle). In contrast, the mouse oocyte has a ratio of approximately 5:1 oocyte to spindle length^18^. The *Drosophila* oocyte itself is vitellogenic and surrounded by protective outer casings, reflecting the need for the oocyte to be as robust as possible as they ultimately develop external to the organism. These features can initially make distinguishing the components of the *Drosophila* spindle challenging. However, with the plethora of genetic tools available, visualization and manipulation of spindle components can now be easily achieved, highlighting *Drosophila* as an important model system for understanding and future research of this field.

Recently, it has been observed that a population of actin exists within the mammalian oocyte that forms a spindle-like structure and has been shown to regulate chromosome alignment and segregation^18^. Treatment of these oocytes with cytochalasin D (cytoD) and knockout of Formin-2, a key nucleator of spindle-like actin in mice, results in the misalignment of chromosomes during metaphase I and chromosome segregation errors during anaphase, often resulting in aneuploidy. Actin was shown to regulate chromosomal movements in part due to control of the kinetochore microtubules (K-fibres), indicating a likely role of actin in microtubule organization more generally^18^. Multi-color 3D-fluorescence microscopy revealed that human oocytes display a population of spindle actin similar to the population in mice, and additionally demonstrated that there is co-localization of γ-tubulin rich minus ends with filamentous actin clusters at the spindle poles^19^. Pharmacological manipulations revealed a co-operation of actin and microtubules at the meiotic spindle, as disruption of the microtubule spindle morphology is directly mirrored by changes to the spindle actin. Taken together, this data suggests that the spatiotemporal organization of actin during oocyte maturation follows microtubule dynamics.

In this study, we utilize advanced imaging in conjunction with pharmacological and genetic manipulation to demonstrate that a population of spindle actin exists in the metaphase I arrested mature *Drosophila* oocyte. We show that the mammalian Formin-2 homologue, Cappuccino (Capu), is required for the formation of the spindle actin network, which undergoes strikingly similar morphological changes to the microtubule spindle during egg activation. Disruption of this network reveals that actin is required for regulating the alignment and segregation of chromosomes during meiosis. Moreover, visualization and manipulation of calcium ions at this transition reveals the importance of calcium signaling for maintaining the morphology of the metaphase spindle and chromosome segregation. Taken together, our data suggests that actin is required upstream of the microtubules to regulate formation of the spindle.

## RESULTS

### Actin is present at the spindle

Mature *Drosophila* oocytes are arrested at metaphase I and held within the ovaries. Ovulation then triggers egg activation during which the metaphase-anaphase transition occurs. The metaphase arrested microtubule spindle can be visualized with the microtubule binding protein Jupiter (Jup) fused to GFP (Jup-GFP) as a comparatively small structure in relation to the rest of the oocyte (Fig. 1a), and lies parallel to the cortex at the dorsal-anterior tip of the oocyte, just below the dorsal appendages. The microtubule spindle forms an elliptical structure with focused poles, with the chromosomes lying centrally in this structure (Fig. 1a, b). It is often the case that the 4th non-exchange chromosomes are visible as a smaller mass at each tip of the main body of chromosomes. Surrounding the oocyte is a layer of follicle cells, which are required during oogenesis for patterning of the oocyte. The follicle cell nuclei are clearly visible in this layer, encompassing the oocyte (Fig. 1a).

**Fig. 1.**
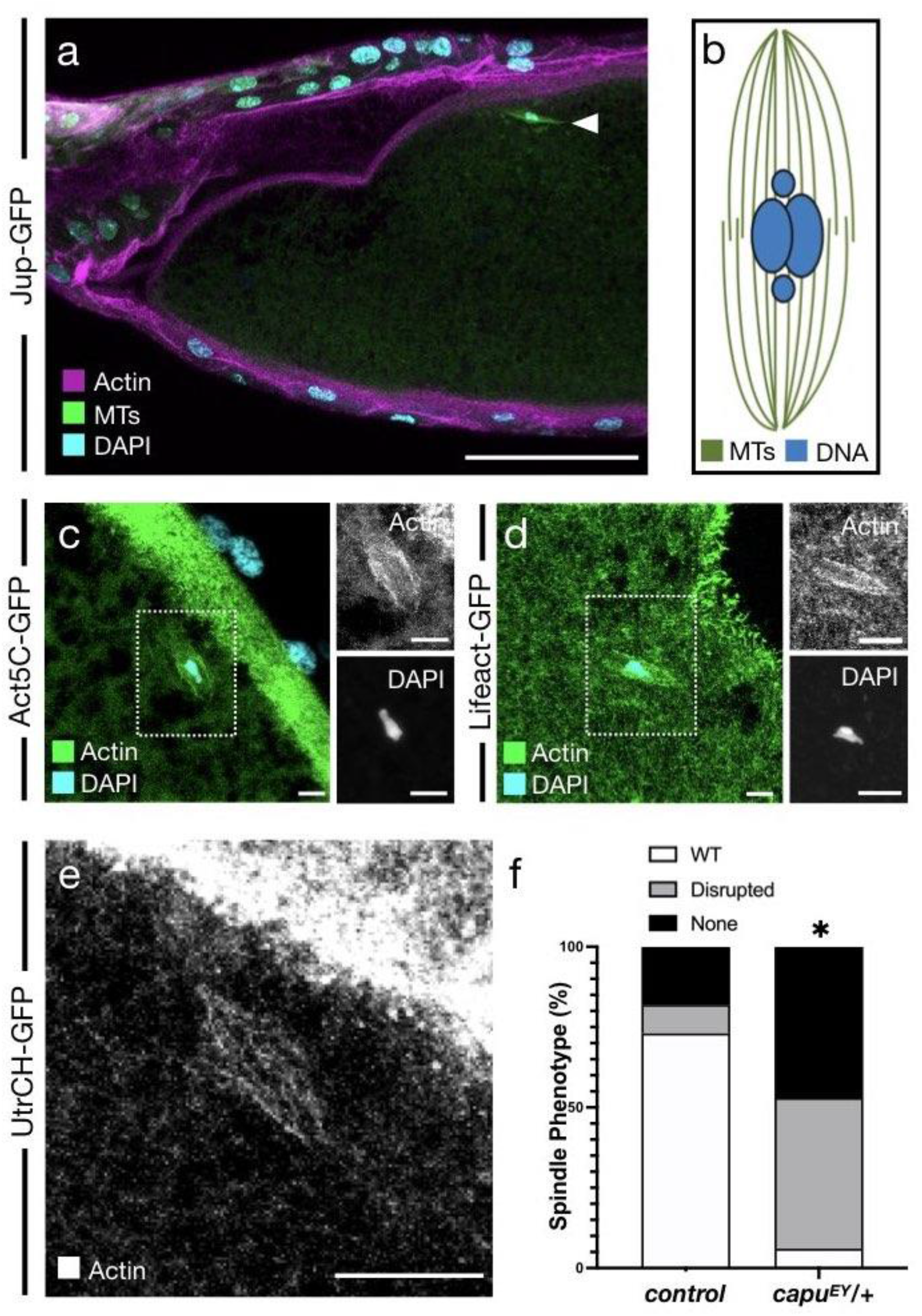
A spindle-like actin population in the metaphase-arrested *Drosophila* mature oocyte. a. Confocal Z-projection (40 μm) of a fixed metaphase I (MI) oocyte. Microtubules shown in green (Jup-GFP, GFP-booster), Actin shown in magenta (Alexa-fluor568 Phalloidin), DNA shown in cyan (DAPI). Spindles (white arrowhead) lie parallel to the dorsal-anterior cortex of the oocyte. b. Schematic representing the MI arrested spindle in the *Drosophila* oocyte. Microtubules (MTs) are represented in green and the metaphase chromosomal mass is represented in blue. Spindle forms with highly focused poles with four centrally located chromosomes. Fourth non-exchange chromosomes appears as a distinct unit at the polar tips of the mass. c. Confocal Z-projection (10 μm) of a fixed MI oocyte. Merge shows actin (Act5C-GFP) in green, DNA (DAPI) in cyan, demonstrating a spindle-like population of actin surrounding a central chromosomal mass. Dashed-line box marks region that is magnified in single-color images to right. d. Confocal Z-projection (10 μm) of a fixed MI oocyte. Merge shows actin (Lifeact-GFP) in green, DNA (DAPI) in cyan, demonstrating a spindle-like population of actin surrounding a central chromosomal mass. Dashed-line box marks region that is magnified in single-color images to right. e. Confocal Z-projection (10 μm) of a live MI oocyte. This population of actin (UtrCH-GFP) appears as a spindle-like structure, with filaments traversing the spindle. f. Comparison of the spindle phenotypes for *wild-type* and *capu*^*EY12344*^*/+* heterozygous mutant shows a significant increase in the percent of oocytes showing elongated or no spindle. N = 25, *<0.05, Fishers Exact Test. Scale bar: 50 μm (a), 5 μm (c,d), 10 μm (e).

We first sought to test if the recently established novel population of actin within the spindle in mouse and human oocytes, is conserved in *Drosophila* oocytes^18,19^. We used two genetically-encoded actin markers: Act5C-GFP, the actin 5C monomer conjugated to a GFP^20^; and Lifeact-GFP, the actin binding protein Lifeact conjugated to a GFP that has been well-established as a faithful actin label in *Drosophila*^21^. These were expressed in the female germ-line, and mature oocytes were fixed and stained with diamidino-2-phenylindole (DAPI) (Fig. 1c, d). A distinct spindle-like population of actin can be observed using both Act5C-GFP and Lifeact-GFP, with the metaphase chromosomal mass localizing to the center of these structures.

This result was also shown live using the calponin-homology domain of Utrophin (Utr), required for the actin binding capacity of Utr, conjugated to an enhanced green fluorescent protein (GFP) (UtrCH-GFP) expressed in the *Drosophila* female germline. This tool has been used previously in other systems to successfully label the spindle actin for live analysis^18^. Utilizing high-resolution confocal microscopy at the dorsal-anterior tip of the oocyte revealed the existence of a small but highly distinct population of actin (Fig. 1e). This actin resembles closely the microtubule spindle as it forms an elliptical shape with focused poles (a unique feature of the *Drosophila* meiotic spindle), and will henceforth be referred to as spindle actin. We then tested whether this population of spindle actin required Capu, the mammalian Formin-2 homologue, for formation. Live visualization of UtrCH-GFP was performed in a *capu* heterozygous background, specifically using *capu*^*EY12344*^, a hypomorphic mutant generated through insertion of a P-element into the first common exon^22-23^. This was sufficient to disrupt the formation of this spindle actin population, with a significant proportion of oocytes not displaying a spindle actin network whatsoever (Fig. 1f).

Together, this establishes a spindle actin population in Drosophila and, consistent with mammals, that the Capu protein is required for accurate formation of the spindle network surrounding the congressed metaphase chromosomes.

### Spindle-like actin regulates metaphase spindle microtubules and chromosomes

To test the function of spindle actin, we first examined the relationship between this actin population and the microtubule spindles. Oocytes expressing Jup-GFP that were incubated in the microtubule depolymerizing agent colchicine showed a complete loss of the microtubule spindle (Fig. 2a). However, the spindle actin (UtrCH-GFP) remained in the presence of colchicine (Fig. 2b), suggesting this population is resistant to disruption of the spindle microtubules. Inversely, depolymerization of the actin cytoskeleton using cytoD resulted in an elongated morphology of the microtubule spindle, as well as the expected loss of spindle actin (Fig. 2c and 4b). To more directly target the spindle actin, we visualized Jup-GFP in mutant backgrounds of *capu* and *spire*, which act as part of a Capu-Spire actin nucleating complex (Fig. 2d-f). In both heterozygous and trans-heterozygous mutant oocytes, the spindle appeared significantly elongated (Fig. 2g). Furthermore, in *capu* trans-heterozygous backgrounds we observe a significant increase in the number of oocytes without spindle microtubules (Fig. 2h).

**Fig. 2.**
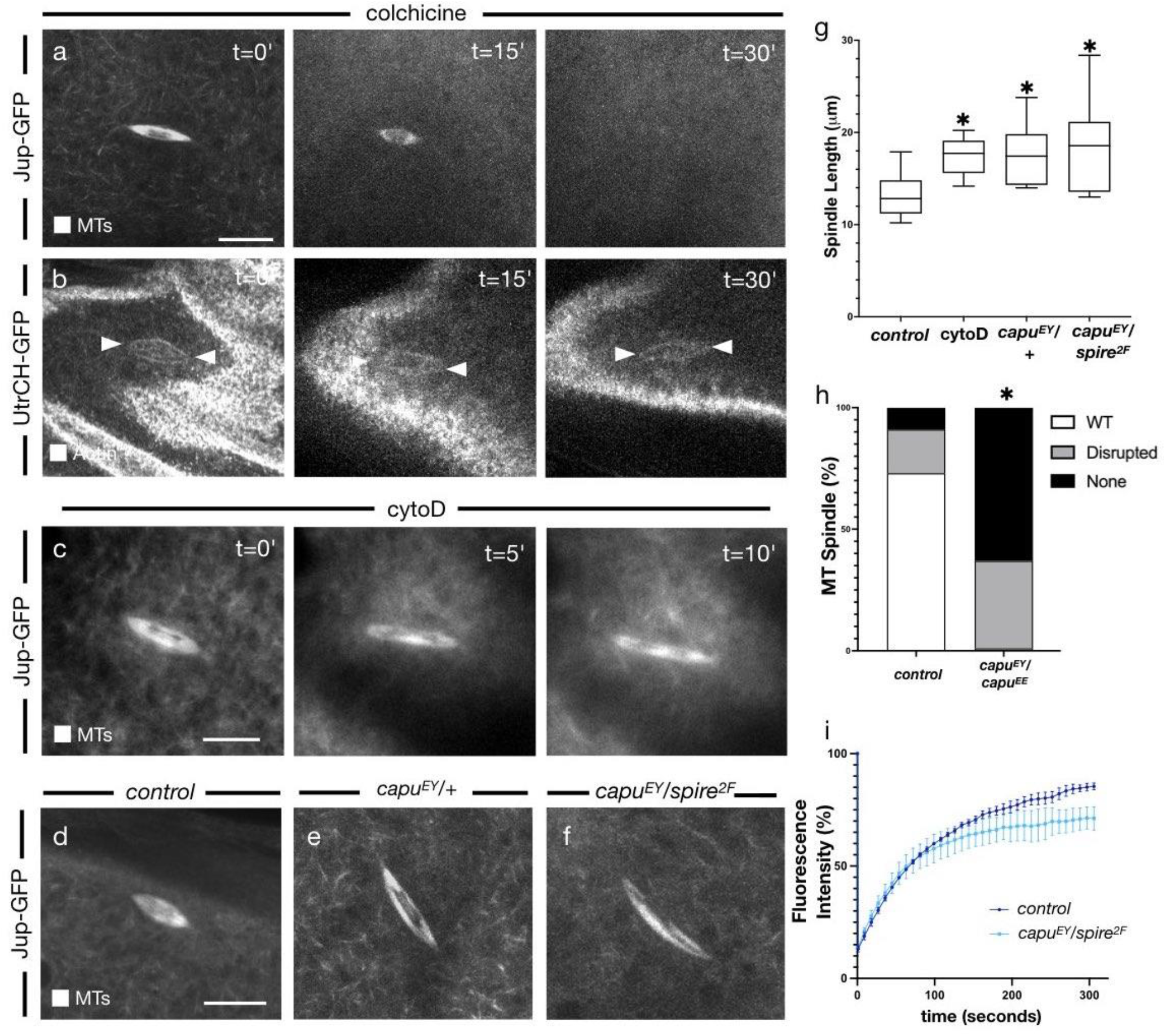
Actin cytoskeleton promotes recruitment of spindle microtubules and regulation of spindle morphology. a. Confocal Z-projections (10 μm) from a live time series of a MI oocyte treated with colchicine (t=0′). Microtubules (Jup-GFP) first appear as a typical spindle structure and completely depolymerize post-treatment (t=30′). N = 5. b. Confocal Z-projections (10 μm) from a live time series of a MI oocyte treated with colchicine (t=0′). Spindle-like actin (UtrCH-GFP) remains unchanged post-treatment (t=30′). N = 5. c. Confocal Z-projections (10 μm) from a live time series of a MI oocyte following incubation with the 10uM of the actin depolymerizing agent cytoD. Before addition (t=0′), the spindle (Jup-GFP) appears as a typical elliptical structure of approximately 10 μM. Post-cytoD addition (t=10′), the spindle has undergone a distinct morphological change as it elongates significantly. Scale bar represents 10 μM. N = 9. d. Confocal Z-projections (10 μm) from a live MI oocyte. Microtubules (Jup-GFP) appear as the typical spindle structure. N = 20. e. Confocal Z-projection (10 μm) from a live *capu*^*EY12344*^*/*+ heterozygous mutant MI oocyte. Microtubules (Jup-GFP) appear as an elongated spindle. N = 20. f. Confocal Z-projection (10 μm) from a live *capu*^*EY12344*^*/spire*^*2F*^ trans-heterozygous mutant MI oocyte. Microtubules (Jup-GFP) appear as an elongated spindle. N = 20. g. Comparison of the microtubule spindle length indicates a significant increase in cytoD treated and mutant backgrounds compared to wild-type. *< 0.05, N = 25, student’s t-test. h. Comparison of the spindle phenotypes for wild-type and *capu*^*EY12344*^*/capu*^*EE*^ trans-heterozygous mutant shows a dramatic increase in the percent of oocytes without a spindle. N = 20, *<0.05. i. Recovery of fluorescence intensity following photobleaching of microtubules in wild-type and *capu*^*EY12344*^*/spire*^*2F*^ trans-heterozygous mutant backgrounds. Mutant oocytes initially show similar recovery dynamics to wild-type oocytes, but overall recovers to a lesser degree. N = 5. Scale bar: 10 μm (a-f).

Next, we used fluorescence recovery after photobleaching (FRAP) to test the dynamics of microtubule recruitment in wild-type and *capu/spire* trans-heterozygous oocytes. When the spindle actin is disrupted, we observed a significant change in the recovery dynamics of the microtubule spindle (Fig. 2i). The failure of the spindle to recover fluorescence to a wild-type level, suggests the spindle actin population plays a role in the recruitment of microtubules to the spindle itself. Taken together, our analyses reveal an important relationship between the spindle actin and microtubule spindle, as actin is required for accurate formation of the microtubule spindle and regulation of its morphology. This appears conserved with studies in mice and humans, in which the spindle actin has been shown to be required for formation of K-fibres and recovery of the spindle structure^18,19^.

Considering the function of the microtubule spindle in chromosome segregation, we next tested if the spindle actin is involved in this process. We, therefore, observed chromosome alignment within the metaphase-arrested spindle after disruption of actin. Prior to metaphase, the chromosomes lie in a centrally congressed mass surrounded by the spindle actin (Fig. 3a), however, following treatment with cytoD (Fig. 3b), there is significant disruption to the alignment of these chromosomes. We detect multiple chromosomal masses spreading to either pole of the spindle axis. Similarly, in c*apu* and s*pire* mutant genetic backgrounds, clear disruption to the centrally congressed chromosomal mass can be observed (Fig. 3c, d). Measuring this spread of the metaphase I chromosomes as the maximum chromosomal distance reveals a significant increase in those oocytes with a disrupted spindle actin (Fig. 3e). Furthermore, analysis of the angle the spindle makes with the cortex reveals a significant change when actin is disrupted (Fig. 3f). Together this suggests that this population of actin is required to promote and maintain the compact nature of the chromosomes centrally, without which misalignment of the chromosomes and spindle occurs.

**Fig. 3.**
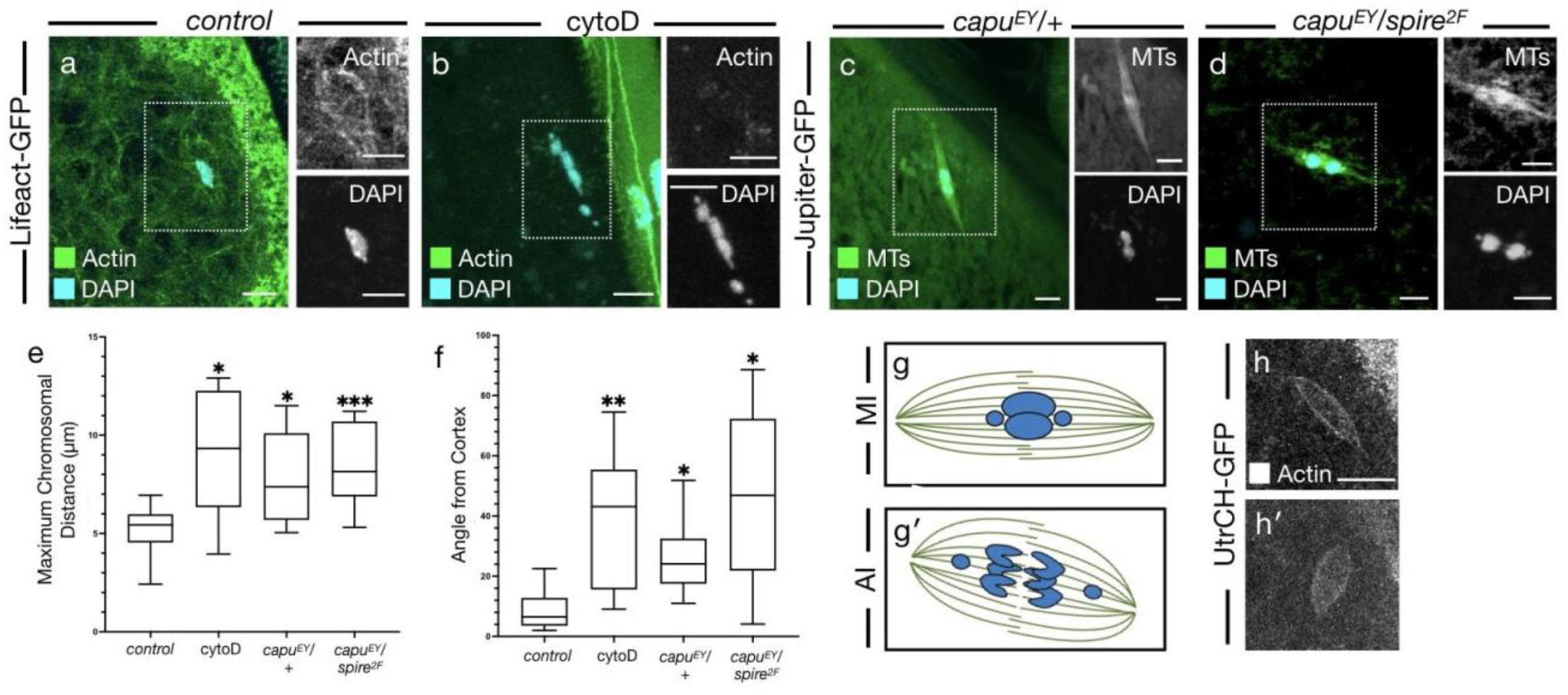
Actin is required for alignment of chromosomes and positioning of the metaphase I spindle. a. Confocal Z-projection (10 μm) of a fixed MI oocyte. Merge shows actin (Lifeact-GFP) in green, DNA (DAPI) in cyan, demonstrating a spindle-like population of actin surrounding a central chromosomal mass. Dashed-line box marks region that is magnified in single-color images to right. b. Confocal Z-projection (10 μm) of a fixed MI oocyte following incubation with the 10uM of the actin depolymerizing agent cytoD. Merge shows actin (Lifeact-GFP) in green, DNA (DAPI) in cyan, demonstrating a loss of the spindle-like population of actin and a drastic separation of the chromosomal mass into several units that elongate along the spindle axis. Dashed-line box marks region that is magnified in single-color images to right. c. Confocal Z-projection (10 μm) of a fixed *capu*^*EY12344*^*/+* heterozygous mutant MI oocyte. Merge shows actin (Jup-GFP) in green, DNA (DAPI) in cyan, demonstrating a separation of the chromosomal mass into two units that begin to migrate along the spindle axis. Dashed-line box marks region that is magnified in single-color images to right. d. Confocal Z-projection (10 μm) of a fixed *capu*^*EY12344*^*/spire*^*2F*^ trans-heterozygous mutant MI oocyte. Merge shows actin (Jup-GFP) in green, DNA (DAPI) in cyan, demonstrating further separation of the chromosomal mass into two distinct units that are migrating to the spindle poles. Dashed-line box marks region that is magnified in single-color images to right. Contrasted to better visualize the spindle. e. Comparison of the maximum chromosomal distance indicates a significant increase in cytoD treated and mutant background compared to wild-type. ***< 0.005, *< 0.05, N = 10, student’s t-test. f. Comparison of the spindle-cortex angle (degrees) indicates a significant increase in cytoD treated and mutant background compared to wild-type. **< 0.01, *< 0.05, N = 10, student’s t-test. g. Schematic representing the MI arrested spindle (g) and anaphase I (AI) spindle (g′) in the *Drosophila* oocyte. Microtubules (MTs) are represented in green and chromosomes are represented in blue. MI arrested spindle is narrow with four centrally located chromosomes (g) whereas the AI spindle is has a greater width, chromosomes separating to opposite poles and is rotating relative to the cortex (g′) h. Confocal Z-projections (10 μm) from a live time series of an MI oocyte pre (h) and post (h′) incubation with activation buffer (AB). Spindle-like actin (UtrCH-GFP) appears narrow pre-AB (h) but undergoes a morphological change as it contracts and widens post-AB (h′). N = 10. Scale bar: 5 μm (a-d), 10 μm (h).

### Functional importance of the spindle actin during anaphase I

In order to test if the functional importance of this actin population extends throughout meiosis, we observed the spindle actin during egg activation. In *Drosophila*, egg activation occurs as the oocyte passes into the oviduct, but can be recapitulated *ex vivo* through incubation in a hypotonic buffer (activation buffer (AB)) which causes the egg to swell. This results in TrpM calcium ion channels opening which enables a calcium transient to pass through the cell and initiate the metaphase-anaphase transition^14,15^. Microtubule spindles undergo a classical rearrangement at egg activation as the spindle initially elongates, then contracts and rotates in relation to the cortex, ultimately becoming perpendicular to the cortex by meiosis II (Fig 3g). Observation of the spindle actin during egg activation reveals a similar morphological change, as the spindle actin ultimately contracts (Fig. 3h). This is highly reminiscent of the morphological change observed with the microtubules at egg activation. This suggests that the interplay between spindle actin and the microtubule spindle continues in anaphase I.

We next tested the role of the actin cytoskeleton during anaphase. Fixation of activated oocytes revealed co-localization of the anaphase I chromosomes with a filamentous and spindle-like population of actin (Fig. 4a). Disruption of the anaphase spindle actin was achieved through treatment with cytoD following AB treatment to ensure oocytes had entered anaphase. This resulted in significant disruption to the segregation of the chromosomes, with frequent occurrence of aberrant chromosomal masses, which we define as chromosomes completely separate from the main axis of segregating chromosomes or a mass causing obvious non-uniformity (Fig. 4b, c). Similarly, visualization of the chromosomes in *capu* heterozygous and c*apu/spire* trans-heterozygous backgrounds revealed aberrations in chromosome segregation (Fig. 4d-g). Individual aberrant chromosomal masses were clearly identifiable, with all test oocytes demonstrating a significant increase in the number of these masses as compared to wild type oocytes (Fig. 4h). Furthermore, in the case of *capu/spire* trans-heterozygous oocytes, it was common to observe the 4th non-exchange chromosomes being closely located, suggesting a loss of spindle polarity (Fig. 4g). Despite clear loss of accurate segregations, measurements of the maximum chromosomal distances did not reveal any significant differences from control oocytes, suggesting chromosome segregation was still able to occur, but with a loss of accuracy (Fig. 4i). In addition, we observed no significant difference in the angle of the spindle-cortex between cytoD treated oocytes in anaphase versus anaphase controls and cytoD treated metaphase oocytes (Fig. 4j). We do, however, still observe a significant increase in the angle as compared to metaphase, with the most significant increase for cytoD treated anaphase oocytes. Together, this demonstrates that many anaphase events are still capable of occurring when the spindle actin is disrupted, including chromosome segregation and spindle rotation. It therefore suggests that the main role of the spindle actin is in providing a level of accuracy to these events, as without it we see dramatic aberrations in the quality of chromosome segregation.

**Fig. 4.**
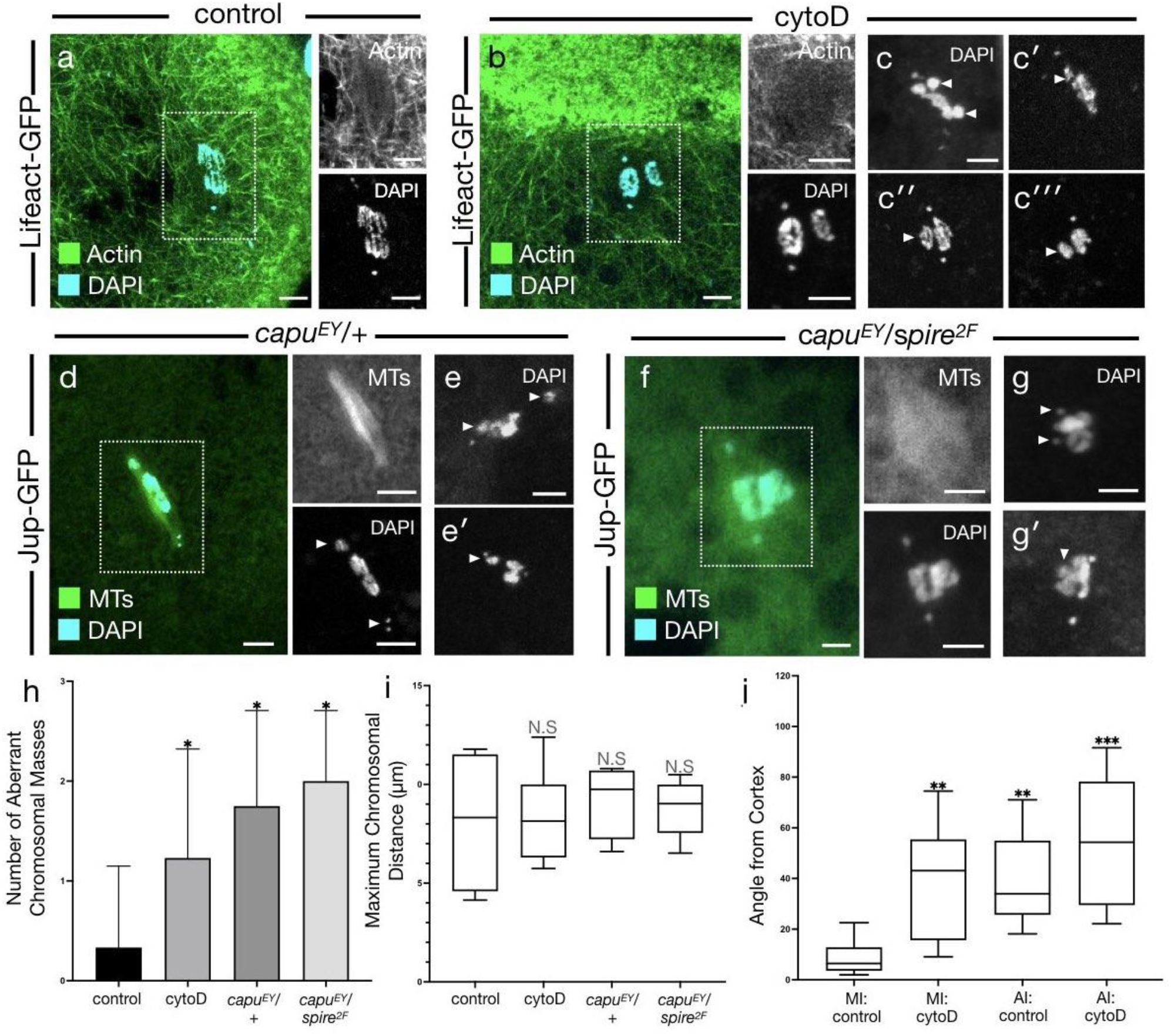
Actin is required for accurate segregation of chromosomes during anaphase I. a. Confocal Z-projection (10 μm) of a fixed AI oocyte. Merge shows actin (Lifeact-GFP) in green, DNA (DAPI) in cyan, demonstrating a filamentous spindle-like population of actin surrounding the segregating anaphase chromosomes. Dashed-line box marks region that is magnified in single-color images to right. N = 15. b. Confocal Z-projection (10 μm) of a fixed AI oocyte following incubation with the 10 μM of the actin depolymerizing agent cytoD. Merge shows actin (Lifeact-GFP) in green, DNA (DAPI) in cyan, demonstrating the loss of a spindle actin population and the disruption of the accurate segregation of the chromosomes. Dashed-line box marks region that is magnified in single-color images to right. N = 10. c. Confocal Z-projections (10 μm) of fixed AI oocytes following incubation with the 10 μM of the actin depolymerizing agent cytoD (C-C′′′). Chromosomes (DAPI) demonstrating further examples of the variety of disrupted chromosome segregation phenotypes, many showing chromosomal units separated from the main mass (white arrowheads). d. Confocal Z-projection (10 μm) of a fixed *capu*^*EY12344*^*/+* heterozygous mutant AI oocyte. Merge shows microtubules (Jup-GFP) in green, DNA (DAPI) in cyan, demonstrating the presence of the spindle but mis-regulation of chromosome segregation. Dashed-line box marks region that is magnified in single-color images to right. N = 12. e. Confocal Z-projections (10 μm) of fixed *capu*^*EY12344*^*/+* heterozygous mutant AI oocytes (e-e′). Chromosomes (DAPI) in cyan demonstrating further examples of the variety of disrupted chromosome segregation phenotypes, many showing chromosomal units separated from the main mass (white arrowheads). f. Confocal Z-projection (10 μm) of a fixed *capu*^*EY12344*^*/spire*^*2F*^ trans-heterozygous mutant AI oocyte. Merge shows microtubules (Jup-GFP) in green, DNA (DAPI) in cyan, demonstrating a reduced microtubule signal and mis-regulation of chromosome segregation. Dashed-line box marks region that is magnified in single-color images to right. N = 8. g. Confocal Z-projections (10 μm) of fixed *capu*^*EY12344*^*/spire*^*2F*^ trans-heterozygous mutant AI oocytes (g-g′). Chromosomes (DAPI) in cyan demonstrating further examples of the variety of disrupted chromosome segregation phenotypes, often showing a complete loss of polarity as the 4th non-exchange chromosomes no longer migrate to poles (white arrowheads). h. Comparison of the number of aberrant chromosomal masses shows a significant increase in cytoD treated and mutant backgrounds compared to the wild-type. *< 0.05, N = 8, student’s t-test. i. Comparison of the maximum chromosomal distance shows no significant difference between cytoD treated and mutant backgrounds. N = 8. j. Comparison of the angle between the spindle and cortex shows a significant increase in cytoD treated and mutant backgrounds compared to the wild-type. There is no significant differences between cytoD and mutant oocytes. **< 0.01, ***< 0.005, N = 8. Scale bar: 5 μm (a-g′)

### Localized calcium signaling may be required at the spindle for effective meiosis

Previous work has shown a clear link between calcium and actin at and around egg activation^24^. Recent evidence in mature *Xenopus* oocytes also highlights the presence of enriched calcium and necessity of localized calcium signaling at the spindle for regulation of microtubules^25^.

We used a genetically encoded calcium sensor, GCaMP3, to visualize calcium *in vivo*^26^. Increased fluorescence intensity at the spindle was observed with live and fixed imaging, indicating a calcium enrichment (Fig. 5a, b). Interestingly, there appears to be the greatest enrichment at the tip of each pole of the spindle in the metaphase oocyte, compared to a more uniform signal in the anaphase oocyte, yet this still suggests a requirement throughout egg activation.

**Fig. 5.**
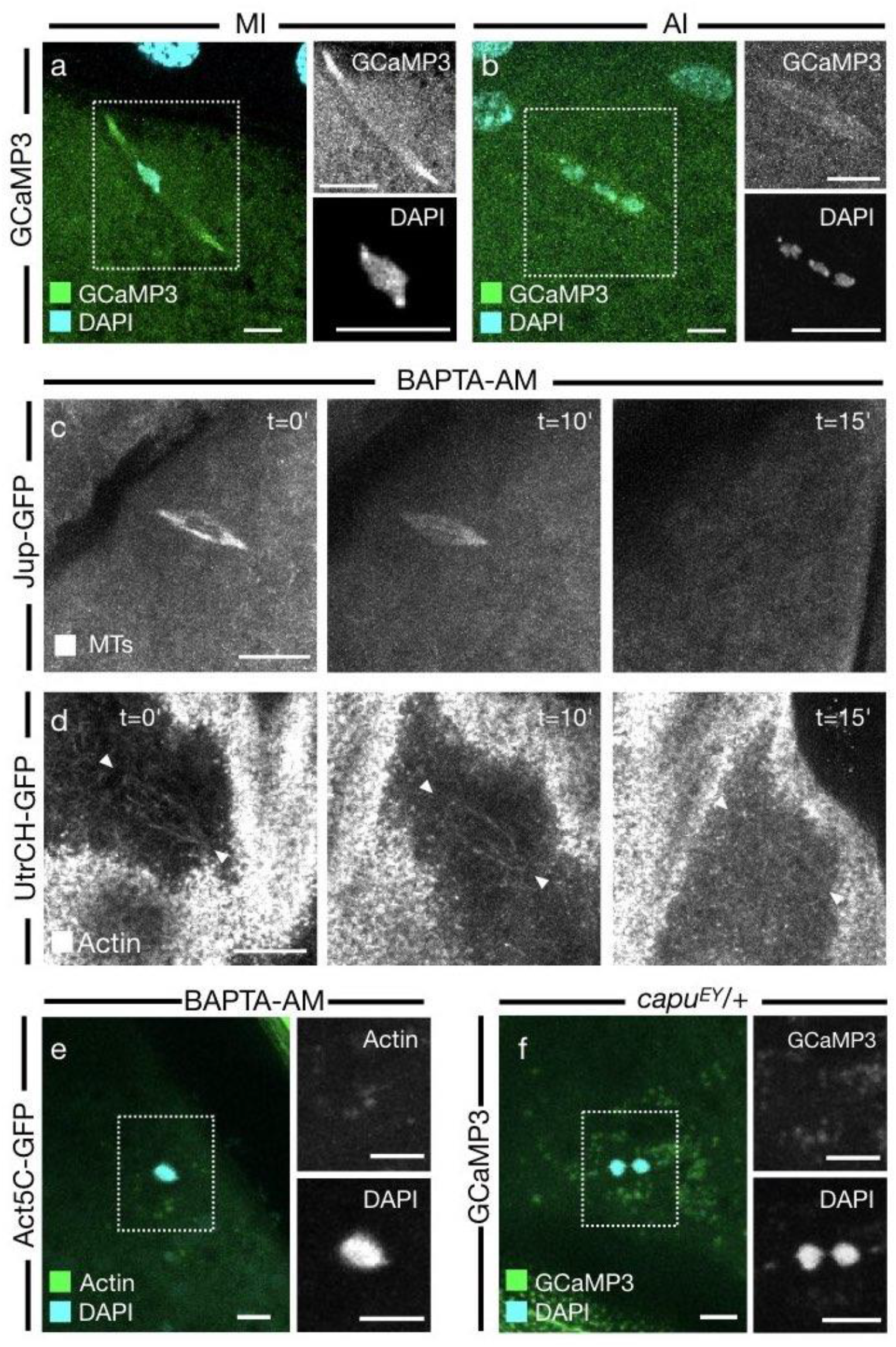
Calcium is required to maintain the metaphase spindle. a. Confocal Z-projection (10 μm) of a fixed MI oocyte. Merge shows calcium (GCaMP3) in green, DNA (DAPI) in cyan, demonstrating a higher calcium signal in the within the spindle. b. Confocal Z-projection (10 μm) of a fixed AI oocyte. Merge shows calcium (GCaMP3) in green, DNA (DAPI) in cyan, demonstrating a separating chromosomal mass and a wider spindle. c. Confocal Z-projections (10 μm) from a live time series of a MI oocyte treated with BAPTA-AM (t=0′). Microtubules (Jup-GFP) first appear as a typical spindle structure and completely depolymerize post-treatment (t=15′). N = 8. d. Confocal Z-projections (10 μm) from a live time series of a MI oocyte treated with BAPTA-AM (t=0′). Actin (UtrCH-GFP) first appear as a typical spindle structure and completely depolymerize post-treatment (t=15′). N = 6. e. Confocal Z-projection (10 μm) of a fixed MI oocyte. Merge shows actin (Act5C-GFP) in green, DNA (DAPI) in cyan, demonstrating a loss of the actin spindle and round chromosomal mass. N = 6. f. Confocal Z-projection (10 μm) of a fixed *capu*^*EY12344*^*/+* heterozygous mutant MI oocyte. Merge shows calcium (GCaMP3) in green, DNA (DAPI) in cyan, demonstrating a loss of the GCaMP3 signal in a spindle conformation and separation of the chromosomes into two separate masses. N = 6. Scale bar: 5 μm (a,b,e,f), 10 μm (c,d).

To test the role of calcium at the spindle, mature oocytes were incubated in the membrane permeable calcium chelating agent BAPTA-AM. The removal of calcium results in the loss of both an actin and microtubule spindle population, suggesting that calcium signaling is required to maintain the spindle apparatus (Fig. 5c, d). This is similar to results in *Xenopus* in which BAPTA incubation caused microtubule depolymerization in a similar fashion to microtubule depolymerizing agents^25^. Furthermore, BAPTA-AM incubations result in the complete compaction of the chromosomes into one rounded mass (Fig. 5e). This phenotype is likely explained by the previous results in which BAPTA causes complete loss of the microtubules and actin within the spindle.

Finally, disruption of the spindle actin through introduction of the capu^EY12344^ mutant resulted in a loss of enrichment of calcium at the spindle (Fig. 5f). As expected, the chromosomes become separated into two individual masses that begin to separate toward spindle poles. However, the enrichment of the GCaMP3 can no longer be observed. Together this shows that calcium is required for the maintenance of the spindle apparatus at metaphase, with an interdependence becoming apparent between the spindle actin and calcium signaling.

## DISCUSSION

This study establishes the existence of a population of spindle-like F-actin in *Drosophila* mature oocytes that is required for the regulation of meiosis. We demonstrate the requirement of the mammalian homologue of Formin-2 in the formation of this spindle actin. Disruption of the actin spindle results in dramatic chromosome segregation errors. At metaphase, the spindle lacks accurate chromosome alignment and congression, and at anaphase aberrant chromosomal masses are frequent. We also identify calcium as critical in maintaining the spindle throughout metaphase. Together, we suggest that the spindle actin is required to mediate the accurate segregation of chromosomes through regulation of the microtubule spindle. Our data, together with recent work that reveals a population of spindle actin in mammals^18,19^ and calcium enrichment in *Xenopus*^27^, suggests there is a high level of evolutionary conservation at the spindle.

The conservation, from mammals to *Drosophila*, we see in both functional and morphological similarities between spindle actin populations. Whilst variation is to be expected in comparison of meiotic mechanisms between species, such as final meiotic arrest occurring at metaphase I in *Drosophila*, in comparison to metaphase II in mammals, it appears that distinct populations of spindle actin may be another fundamental feature of meiosis. The conservation of this population of actin extends from its nucleation by the Formin-2 homolog Capu, to its functional role in the regulation of spindle microtubules and chromosomal movements.

However, there does appear to be some variation in the morphology and function of this spindle population. Much like the microtubule spindle, the spindle actin forms an elliptical shape with highly focused poles, unlike mammalian meiotic spindle structures which are more ‘barrel like’ in shape^2^. This population also seems to be resilient to disruption of the microtubule spindle, appearing to be important for recruitment of microtubules to the spindle and regulating overall spindle morphology, therefore suggesting an upstream requirement of the actin cytoskeleton. When disrupted, either through knockdown of *capu* or depolymerization by cytoD, significant defects in chromosome alignment can be observed. Observation of metaphase I oocytes indicated a spreading of the chromosomes along the metaphase spindle, with separation of chromosomes appearing reminiscent of pro-metaphase oocytes^17^. This could suggest a requirement of the spindle actin in the prophase-metaphase transition, a much earlier stage than has been observed in mammals to date. Perhaps this population of actin plays a more significant role in *Drosophila* meiosis I as mature oocytes arrest at metaphase I, which may indicate that the actin is important in meiotic arrest and release from this arrest during egg activation and onset of the metaphase-anaphase transition.

During egg activation, a global transient of calcium triggers a plethora of events, including global rearrangements of the actin cytoskeleton and resumption of meiosis^24^. There is building evidence that ties calcium and actin as two interlinked molecules in many signaling pathways; in *Drosophila* egg activation, dispersion of cortical actin enables entry of calcium in the form of a wave, which, downstream, effects a wave of reorganizing F-actin^24^. With many actin binding proteins (ABPs) being calcium sensitive, such as α-actinin and the villin family, and many calcium-sensitive proteins having downstream effects on the actin cytoskeleton, such as Calmodulin and calcineurin^28^, it is very probable that the calcium wave has a direct effect on the population of spindle actin, potentiating the release from meiotic arrest.

We have observed directly an enrichment of the calcium indicator GCaMP3 at the metaphase arrested spindle, suggesting an increased local concentration of calcium. When calcium is removed, depolymerization of the actin and microtubule spindles can be observed, indicating the requirement of calcium in maintenance of these populations, consistent with observations in *Xenopus*^25^. Our data suggests an interdependent relationship with the spindle actin and calcium, as it appears that enrichment of the calcium signal is reduced in *capu* mutants. This may suggest the presence of localized calcium signaling between the spindle actin and associated ABPs and/or that the actin aids in recruitment of calcium-sensitive proteins more generally within the spindle. More complete knockdown of c*apu* often results in the loss of formation of the microtubule spindle, furthering an argument in which the spindle actin is required for recruitment of microtubules. This recruitment could be direct, but given the requirement of calcium for formation and maintenance of spindle populations, it is likely that the loss of actin may impact the formation of microtubules through disruption of the localized action of many calcium sensitive proteins. One family of candidates is the microtubule associated proteins (MAPs), many of which are calcium sensitive and associate with both the actin and microtubule cytoskeleton^27^. Without the spindle actin, it is likely that the localization and coordination of these proteins are lost, resulting in a loss of calcium signaling at the spindle and concomitant loss of the microtubule spindle itself. Given the ubiquitousness of calcium at the spindle, this mechanism should be explored further in mammals, as it is likely another conserved feature of meiosis.

## ACKNOWLEDGEMENTS

We are grateful to Jens Januschke, Torsten Krude, Jose Casal, and Paul Conduit for discussions and advice; Elise Wilby for feedback on the manuscript; the Zoology Imaging Facility and Matt Wayland for assistance with microscopy; the Bloomington *Drosophila* Stock Center and *Drosophila* community for fly lines.

## AUTHOR CONTRIBUTIONS

B.W.W. was responsible for conceptualization, methodology, experimentation, analysis, and writing. T.T.W was responsible for conceptualization, supervision, and reviewing.

## MATERIALS AND METHODS

### Fly Maintenance

Fly stocks were raised on Iberian recipe fly food at 18°C, 21°C and 25°C. For dissection of mature oocytes, approximately 30 female flies with 5 male flies were transferred into a vial with Iberian recipe fly food and wet yeast for 48 hours at 25°C.

### Fly lines

*matα-GAL4::VP16, UASp-GCaMP3* ^29^ ; *tub-GAL4VP16* (S. Roth); *UASp-Utrophin-CH-GFP/Tm3* ^30^; *UASp-LifeactGFP/Tm3* (BL58717); *UASp-Act5CGFP/Tm3* (BL7309); *P{PTT-GA}JupiterG00147* (BL6836); *P{PTT-un}CamP00695*/CyO (BL50843); *capu*^*EY12344* 30^; *capu*^*EE*^*/CyO* (BL8788); *spire*^*2F*^*/CyO* (BL8723).

### Preparing oocytes for live imaging

Ovaries from flies fattened for 48 hours on yeast were dissected onto a 22 by 40 mm cover slip, into series 95 halocarbon oil using forceps (11251-30 Dumont #5 forceps, Fine Science Tools) and a dissecting probe (0.25 mm straight 10140-01, Fine Science Tools) as described previously^31^. Mature oocytes were gently teased out of the ovaries and left for 10 minutes prior to imaging to allow them to settle onto the coverslip.

### Preparing *in vivo* activated eggs

Ovaries from flies fattened for 48 hours on yeast were carefully dissected onto a 22 by 40mm cover slip, such that the oviduct was still intact, into series 95 halocarbon oil. Oocytes were then gently teased out of the oviduct, carefully removing the surrounding oviduct tissues.

### Preparation of fixed samples

Ten to twenty ovaries were dissected from flies fattened for 48 hours on yeast into Schneider’s Insect Medium (Sch) (GibCo). Ovaries were splayed open and oocytes gently teased out using fine forceps (11251-30 Dumont #5 forceps, Fine Science Tools) and a dissecting probe (0.25 mm straight 10140-01, Fine Science Tools). Mature oocytes were transferred into an 0.5mL eppendorf tube using a glass pipette. Sch was removed and 500 μL 4% paraformaldehyde (PFA) stabilized with phosphate buffer (Thermofisher Scientific) was added for 10-15 minutes on a rotary machine (PTR-35 360 vertical multi-function rotator, Thermofisher Scientific). Oocytes were then washed for 10 minutes, three times in 0.1% PBST (0.1% Triton X-100 (ThermoFisher Scientific) in PBS. Oocytes were then incubated for 2 hours in 1% PBST with the following labelling probes, before washing, staining in glycerol and DAPI and mounting on a glass slide in Vectashield with DAPI (Vector Laboratories): Alexa-Fluor Phalloidin 568, 1:500 (Molecular Probes); Alexa-Fluor Phalloidin 637, 1:500 (Molecular Probes); ChromoTek GFP-booster, 1:500 (Proteintech).

### *Ex vivo* egg activation

Mature oocytes were activated *ex vivo* through addition of the hypotonic, 260 mOsm, activation buffer (AB): 3.3 mM NaH_2_PO_4_, 16.6 mM KH_2_PO_4_, 10mM NaCl, 50 mM KCl, 5% polyethylene glycol (PEG) 8000, 2mM CaCl_2_, brought to pH 6.4 with a 1:5 ratio of NaOH:KOH ^32^. Oocytes typically activate within two minutes of addition of AB.

### Pharmacological treatments

All pharmacological incubations were validated with either a Schneider’s Insect Medium (Sch) or PBS control and a DMSO control made to the appropriate dilution. Oocytes were incubated in Sch or PBS for 10 minutes whereupon samples were flooded with AB and visualized.

CytoD (Sigma Aldrich) was made to a final concentration of 2-20 μM in Sch or PBS. Oocytes were dissected into Sch or PBS a glass bottom dish. The Sch or PBS was carefully removed and replaced with the cytoD. Oocytes were incubated for 10 to 30 minutes in this solution prior to fixation or live imaging. Colchicine (Sigma Aldrich) was made to a final concentration of 50 μM in Sch or PBS. Oocytes were incubated for at least 30 minutes in this solution prior to fixation or live imaging, as above.

### Imaging with the Inverted Olympus FV3000 system

An 1.05 NA 30X silicone objective was used for whole oocyte imaging and an 1.35 NA 60X silicone objective for visualizing intracellular components. For high resolution imaging of the spindle, oocytes were oriented on the coverslip such that the dorsal appendages were in contact with the surface of the cover slip, therefore the dorsal side of the oocyte becomes the shallowest plane of visualization. Parameters for image collection were: 1.35 NA 60x silicon immersion objective, 10 μm Z-stack, 0.5 μm between each Z-slice, 1024×1024 pixels, approximately 15 seconds per stack.

### FRAP analysis

For FRAP of spindle components Jup-GFP was bleached for 10 seconds. Time lapse series of recovery was recorded every 5 seconds in single plane imaging of the cortex or every 30 seconds in Z-stack imaging of the spindle, both using the 488nm laser channel, 2 Airy unit pinhole, 1024×1024 pixels. For all FRAP series, background correction was performed by subtracting the fluorescence intensities of the unbleached cytoplasmic area from fluorescent intensities of bleached regions, with percentage fluorescence of the maximum plotted in graphs.

